# Protein dynamics enables phosphorylation of buried residues in Cdk2/Cyclin A-bound p27

**DOI:** 10.1101/2020.03.12.989517

**Authors:** João Henriques, Kresten Lindorff-Larsen

**Affiliations:** Structural Biology and NMR Laboratory & Linderstrøm-Lang Centre for Protein Science, Department of Biology, University of Copenhagen, Denmark

## Abstract

Proteins carry out a wide range of functions that are tightly regulated in space and time. Protein phosphorylation is the most common post-translation modification of proteins and plays key roles in the regulation of many biological processes. The finding that many phosphorylated residues are not solvent exposed in the unphosphorylated state opens several questions for understanding the mechanism that underlies phosphorylation and how phosphorylation may affect protein structures. First, since kinases need access to the phosphorylated residue, how do such buried residues become modified? Second, once phosphorylated, what are the structural effects of phosphorylation of buried residues and do they lead to changed conformational dynamics. We have used the ternary complex between p27, Cdk2 and Cyclin A to study these questions using enhanced sampling molecular dynamics simulations. In line with previous NMR and single-molecule fluorescence experiments we observe transient exposure of Tyr88 in p27, even in its unphosphorylated state. Once Tyr88 is phosphorylated, we observe a coupling to a second site, thus making Tyr74 more easily exposed, and thereby the target for a second phosphorylation step. Our observations provide atomic details on how protein dynamics plays a role in modulating multi-site phosphorylation in p27, thus supplementing previous experimental observations. More generally, we discuss how the observed phenomenon of transient exposure of buried residues may play a more general role in regulating protein function.

**Significance Statement:** Protein phosphorylation is a common post-translation modification and is carried out by kinases. While many phosphorylation sites are located in disordered regions of proteins or in loops, a surprisingly large number of modification sites are buried inside folded domains. This observation led us to ask the question of how kinases gain access to such buried residues. We used the complex between p27, a regulator of cell cycle progression, and Cyclin-dependent kinase 2/Cyclin A to study this problem. We hypothesized that transient exposure of buried tyrosines in p27 to the solvent would make them accessible to kinases, explaining how buried residues get modified. We provide an atomic-level description of these dynamic processes revealing how protein dynamics plays a role in regulation.

## Introduction

p27^*Kip*1^ (p27) is an intrinsically disordered protein (IDP) that folds upon binding to cyclin-dependent kinase (Cdk)/cyclin complexes, inhibiting their kinase activity (***Polyak et al., 1994*; *Toyoshima and Hunter, 1994; Russo et al., 1996; Galea et al., 2008; Tsytlonok et al., 2020***). These complexes control cell division (***Morgan, 1997***) and thus p27 plays a central role in cell-cycle regulation. The level of p27 protein expression is known to decrease during tumor development and progression, and p27 is a prognostic factor in various human cancers (***Lloyd et al., 1999***). Relief of p27’s inhibition of Cdk can be achieved through phosphorylation of p27 on one or two tyrosine residues — Y88 or Y88 and Y74 — by the non-receptor tyrosine kinases (NRTKs) BCR-ABL and Src, respectively (***Grimmler et al., 2007; Chu et al., 2007***). The resulting partial reactivation of the Cdk/cyclin complex then leads to the phosphorylation of T187 in p27 by Cdk itself, followed by ubiquitination and degradation of p27, thus triggering the full activation of the Cdk/cyclin complex and progression to the next phase of the cell cycle (***Grimmler et al., 2007; Galea et al., 2008***).

While the overall mechanism of the regulation of Cdk/cyclin by phosphorylation of p27 is relatively well understood, key questions remain at the molecular level. In particular, the key regulatory modification sites, Y88 and Y74, are both inaccessible for enzymatic modification in the ternary complex, with Y88 in particular being deeply buried in the active site of Cdk (Fig. 1) (***Russo et al., 1996***). This raises the important question regarding how these sites become accessible for enzymatic modification in signal transduction. Interestingly, a significant number of function-altering phosphorylation sites across the proteome are presumed to be kinase inaccessible in the unphosphorylated state (***Vandermarliere and Martens, 2013; Sirota et al., 2015***).

**Figure 1.**
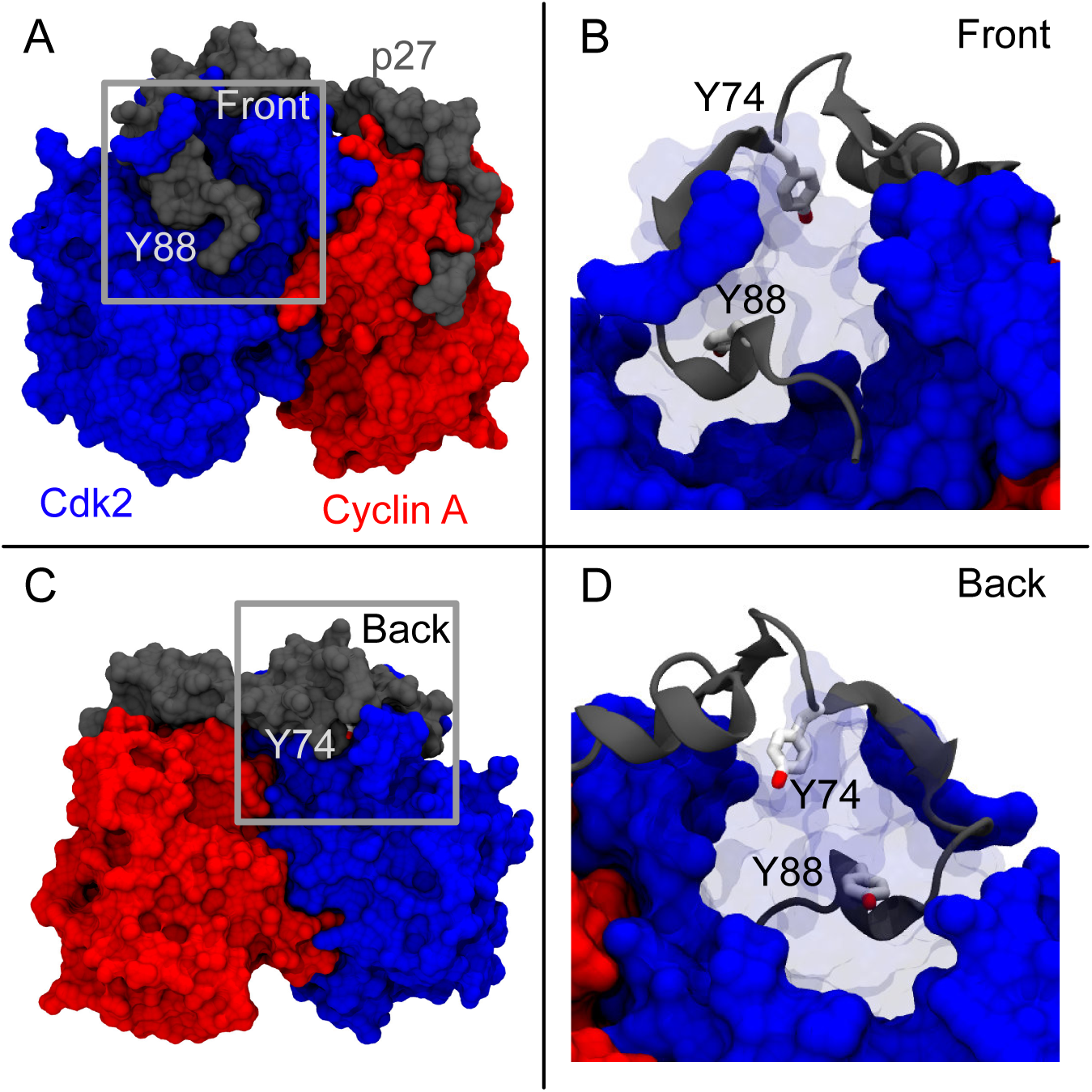
Schematic representation of the ternary complex between p27/Cdk2/cyclin A (PDB 1JSU) (***Russo et al., 1996***). The complex is shown from two different angles (front view (a-b); back view (c-d)). (b) and (d) are detailed representations of the areas corresponding to the gray squares in (a) and (c), respectively. Regions of Cdk2 have been made transparent to highlight the buried tyrosine residues.

One possible mechanism for how kinases gain access to buried substrates has been studied in the context of the Dbl homology (DH) domain of the proto-oncoprotein and guanine nucleotide exchange factor Vav1. Vav1 is autoinhibited through interactions between its catalytic site and a helix from an N-terminal acidic region (***Aghazadeh et al., 2000***) which includes Y174. In analogy to p27/Cdk/cyclin, Y174 is completely buried in the helix-DH interface, yet its phosphorylation is responsible for relieving Vav1 autoinhibition (***Aghazadeh et al., 2000***). Through a combination of NMR and mutagenesis experiments it was shown that Y174 in Vav1 is transiently exposed, and that the rate of phosphorylation correlates with the population of this kinase-accessible state in a series of mutant variants(***Li et al., 2008***). It is thus likely that the presence and population of such an exposed state must be tightly controlled because too low a population would prohibit kinase-accessibility and ability to be phosphorylated, and too high a population would prevent autoinhibition. These important findings motivated the present work, as we hypothesized that a similar dynamic equilibrium between a dominant buried state and an transiently open, kinase-accessible state could be present in the p27/Cdk/cyclin ternary complex, and might — on a broader scope — be a general mechanism involved in protein regulation.

The hypothesis that dynamics and transient exposure of phosphosites in p27 plays a central role in its regulation was recently studied through NMR and single-molecule fluorescence spectroscopy, as well as through biochemical experiments, by showing that Cdk2/Cyclin A-bound p27 samples low populated conformations that dynamically anticipate the sequential steps of the signaling cascade (***Tsytlonok et al., 2019***). The authors termed this ‘dynamic structural anticipation’ (***Tsytlonok et al., 2019***) and showed that even when it is bound to Cdk2/Cyclin A, p27 is intrinsically dynamic across broad timescales (≥ 1.7 ms) (***Tsytlonok et al., 2019***). Nevertheless, a detailed picture of the key interactions governing this equilibrium remains elusive, along with the mechanism behind it. Here, we used enhanced sampling, atomistic molecular dynamics (MD) simulations to show that state-of-the-art physics-based protein models (force fields) can not only capture the aforementioned ‘dynamic structural anticipation’, but also provide detailed information about it that complements experimental observations. We find that in addition to the native contacts between Y88 in p27 and the Cdk2 active site, the contacts of its highly conserved neighbouring residue, F87, are also important. These residues occupy the ATP-binding site of Cdk2 in the unphosphorylated complex and bind to and leave this restricted space it in a cooperative fashion. Additionally, polar interactions between the backbone carbonyl groups of F87 and R90 (within p27) with the highly conserved K33 in Cdk2 are shown to be important for the stability of the bound state, functioning as a latch which helps to keep a 3_10_-helix in p27 in place, effectively sealing the Cdk2 active site. Upon phosphorylation of Y88, we find that the otherwise minor state associated with the transient solvent exposure of Y88, is now spontaneously populated in unbiased molecular dynamics simulations. Further, we examine the transient exposure of Y74 in p27 and how it depends on the phosphorylation status of Y88, and suggest a potential mechanism of coupling between these sites. Together with previous experimental observations our simulations provide an atomic-level description of how transient exposure of buried tyrosine residues occurs and how these processes play a role in the regulation of cell cycle.

## Results and Discussion

### Phosphorylation sites in p27 remain buried in microsecond-length MD simulations

We first performed two standard (unbiased) MD simulations of the ternary complex of p27/Cdk2/Cyclin A in its unphosphorylated state (simulations A and B in Table S1) to test the stability of this complex in the force fields employed, and to examine whether enhanced sampling techniques would be required to study the hypothesized intrinsic dynamics of bound p27. We selected two state-of-the-art force fields from different families and, after an initial relaxation period, no major structural changes were observed (Fig. S1). To monitor the interactions between the two important tyrosines in p27 (Y74 and Y88) and Cdk2 we designed collective variables (CVs) to be sensitive to not only the specific positioning of these residues in relation to the crystallographic structure, but also to all their interactions with C_*α*_ and side-chain carbon atoms of nearby residues. Analyses of these unbiased simulations show that, on this timescale, the phosphorylation sites within the D2 domain of p27 remain fully buried as in the original crystal structure. We did observe small fluctuations in the number of contacts between Y88 and Y74 and the remainder of the complex (CVs VI and XI, Tables S2 and S3) in the beginning of these simulations (Fig. S1,≈ 30 − 40 ns), and attribute these to side chain reorientations as a result of the initial relaxation of the ternary complex. As such, once the initial equilibration is overcome, the coordination numbers quickly stabilize at values similar to those obtained in the crystal structure (Fig. S1).

### Metadynamics simulations show an equilibrium between buried and transiently exposed Y88

Experiments show that oncogenic non-receptor tyrosine kinases BCR-ABL and Src are both able to phosphorylate Y88 in Cdk2/cyclin A-bound p27 (***Grimmler et al., 2007; Chu et al., 2007***). This post-translational modification alone is responsible for the partial reactivation of Cdk2 activity and is generally considered to constitute the first step in the downstream signalling that culminates with the full reactivation of Cdk2 and degradation of p27 (***Tsytlonok et al., 2019***). In addition to Y88, the oncogenic kinase Src can also phosphorylate Cdk2/Cyclin A-bound p27 at Y74. Singly Y74-phosphorylated p27 has not been observed in cells (***Tsytlonok et al., 2019***), and despite the heightened Cdk2 activity associated with the dual phosphorylation of Y88 and Y74, the latter appears to play an ancillary role. It is for these reasons that we first focus on Y88.

We anticipated that any dynamical processes that would lead to the transient exposure of Y88 (and Y74) would take place on timescales beyond those easily accessible to standard MD simulations, and indeed this hypothesis was recently confirmed by experiments that put the timescales at ≈ 1.7 ms or longer (***Tsytlonok et al., 2019***). Thus, we turned to metadynamics as an enhanced sampling simulation technique designed to accelerate the sampling of rare events (***Laio and Parrinello, 2002***). Metadynamics uses a history-dependent biasing potential to enhance simulations and estimate free-energy profiles. In order to be efficient, however, such simulations requires the input of pre-speci1ed collective variables that capture key aspects of the slow dynamics. We have previously used such simulations to sample slow processes in model proteins (***Papaleo et al., 2014; Wang et al., 2016***) where the end points of the conformational exchange were known, and thus in those studies we used that information to design the relevant CVs. In the case of p27 only the starting structure is known, and thus we designed a set of key CVs with the idea that at least some of these would enhance the sampling of the unbinding and rebinding of Y88. After an initial exploratory set of simulations, we selected a number of these for use in more extensive sampling (Table S2). Because standard metadynamics is difficult to converge when using many CVs, we instead used a parallel-bias approach in which we simultaneously apply multiple low-dimensional bias potentials.

In our single walker parallel-bias metadynamics simulation we observe reversible release and rebinding of Y88 from the ATP-binding pocket of Cdk2, as monitored for example by the number of contacts formed between Y88 and its binding site (Fig. 2a, upper panel). The simulations thus reveal both how Y88 gets exposed for putative action of NRTKs (Supporting Movie 1) and subsequently bringing the whole ternary complex back to its dominant state, consistent with the starting X-ray structure (Supporting Movie 2). Solvent exposure of Y88 is achieved through the release of the C-terminal region of p27, roughly corresponding to the D2.3 subdomain (***Sivakolundu et al., 2005***), i.e., residues ≈ 83 − 93 (Fig. 2c, green), in good agreement with single-molecule fluorescence anisotropy (smFA) experiments (***Tsytlonok et al., 2019***). We did not find other major conformational changes in p27 and, importantly, Y74 remains buried during these transient exposures of Y88.

**Figure 2.**
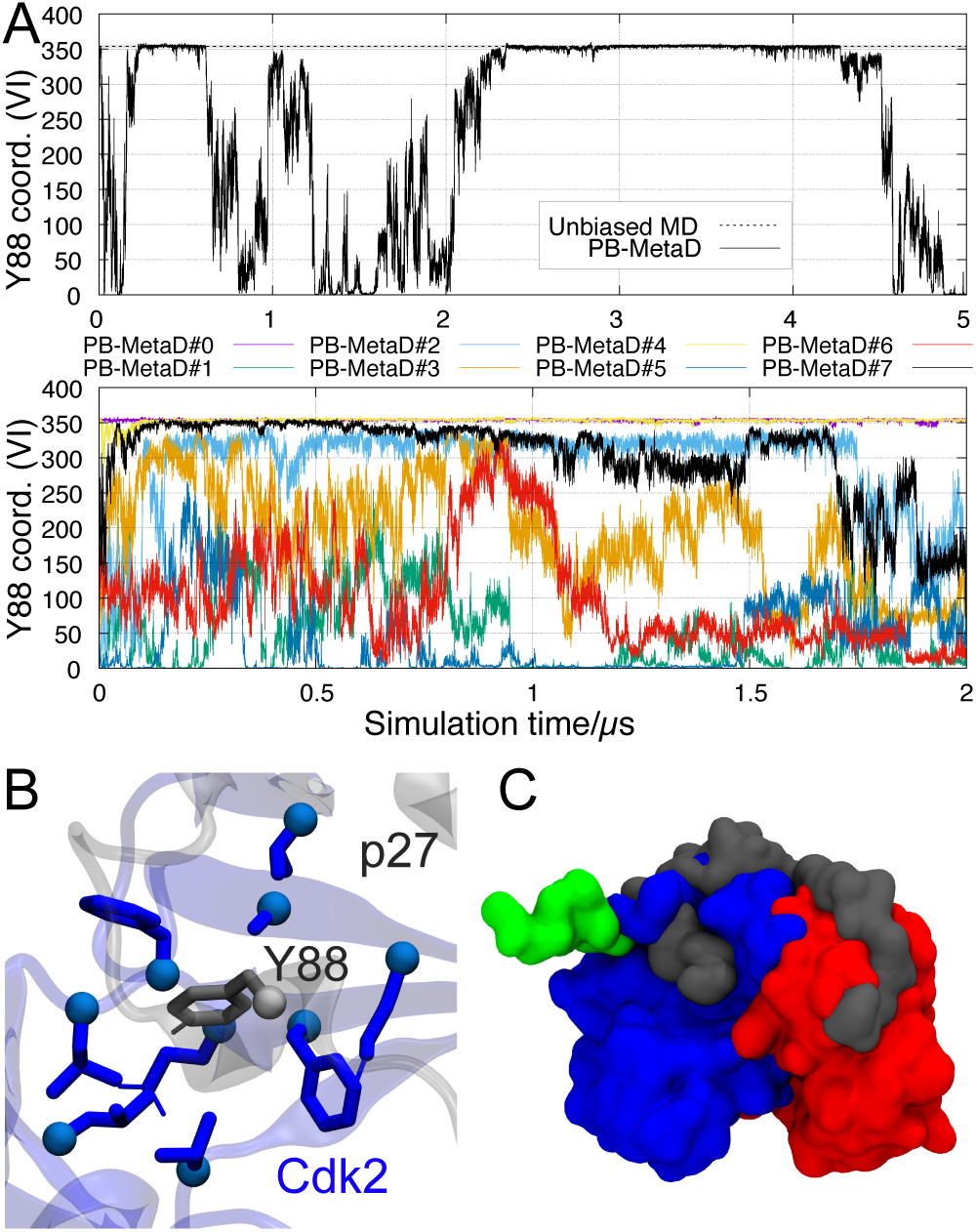
Transient exposure of Y88 in metadynamics simulations. (a) Time series of a CV that quantifies the number of contacts between Y88 and Cdk2 (CV VI, Table S2) in (upper panel) our single-walker parallel bias metadynamics simulation (Table S1, ref. C) and (lower panel) a multiple walker setup with 8 replicas (Table S1, Ref. D). The average number of contacts calculated from our unbiased MD simulation is shown with a dashed line. (b) The number of contacts between Y88 (in gray) and its neighboring residues in Cdk2 (blue) is a good descriptor of the extent to which Y88 is buried in the active site or solvent exposed. (c) Representative structures of a bound, native-like conformation of p27 structure with a high coordination numbers (gray), and a conformation with a low coordination number in which Y88 and the entire D2.3 domain (green) (***Sivakolundu et al., 2005***) is unbound from Cdk2 active site (blue). No other major conformational changes are observed in bound p27. Time series for other CVs that we biased in the multi-replica simulations are shown in Fig. S2. In this and other figures, a roman numeral in a parenthesis refers to the entries in Tables S2 and S3.

While this simulation provides useful detail about the binding/unbinding mechanism of Y88, we only observe two rebinding events to native conformations making it unclear how robustly mechanistic and energetic conclusions may be drawn. The observation that we can reversibly sample the bound and unbound states in microsecond-length enhanced sampling simulations, however, suggest that the CVs capture key aspects of the slowest motions. Thus, we used the same set of CVs in a multi-replica simulation (‘multiple-walkers metadynamics’), where each semi-independent simulation shares and contributes to the same history-dependent bias potential. We obtain additional sampling of the dynamic exchange between buried, partially buried and fully solvent exposed Y88 with the use of eight walkers starting from different points in CV space (Fig. 2a (lower panel) and Fig. S2). Nevertheless, although each simulation is 2 µs long (i.e. an aggregate of 16 µs) we are still not able to produce an unequivocally converged simulation. We base this argument on the fact that we only observed a small number of transitions between the native-like bound state (Y88 coordination number between ≈ 345 − 360) and closely related intermediate states (Y88 coordination number between ≈ 300 − 340). In fact, these intermediate states appear to produce a divide whereby walkers sampling conformations below this value are ‘kept’ on this side of CV space and walkers #0 and #3 seem unable to follow the opposite direction, i.e. unbind. Walker #7 appears to be the exception, and crosses this region on two occasions, i.e. unbound → bound → unbound (≈ 0.15 − 0.55µs). We did not observe this behaviour in the single walker counterpart (Fig. 2a, upper panel).

To examine the transition between this apparent intermediate state and the fully bound state we initiated an unbiased MD simulation from the intermediate state (Table S1, ref. E) and observe that p27 spontaneously progresses from the intermediate state into a native-like conformation along what appears to be similar to the lowest free-energy path (Fig. S3). Thus, in spite of the difficulty in converging the simulations, it is clear that an equilibrium between buried and exposed Y88 states exists and that the simulations provide us with an opportunity to examine the key interactions governing it. Of the eight collective variables we chose to bias in order to promote this dynamic exchange, we observe that three of them (CVs II, III and V, Table S2) correlate with the coordination number of Y88 (CV VI, Table S2), and that, along with visual inference of the sampled binding and unbinding events, suggests that these interactions play a role in controlling the so-called dynamic structural anticipation responsible for making Y88 available for phosphorylation.

In the crystal structure of the ternary complex of p27/Cdk2/Cyclin A, p27 residues F87 and Y88 are responsible for occupying the active site of Cdk2 (Fig. 3d), making it fully inaccessible to ATP. These residues are also highly conserved in p27 homolgues in 65 different species (Fig. S4), and, in fact, F87 is more commonly found than, for example, Y74, suggesting it is of central importance for p27 function. Our metadynamics simulations (Table S1, refs. C and D) reveal a cooperative process, in which these residues access/exit the active site one after the other in a stepwise fashion (Fig. 3a). specifically, examining the two-dimensional free-energy landscape we observe an intermediate state that appears to be on pathway to the fully bound state in which Y88, but not F87, is buried (Supporting Movies 1 and 2). As our simulations are not fully converged we hesitate in making strong quantitative statements, but note that if these variables are used to estimate the free-energy difference between bound and exposed states we find a value (≈ 1.7 kcal/mol) relatively similar to experiments(***Tsytlonok et al., 2019***) (≈ 0.4 kcal/mol). In addition to the interactions made between F87 and Y88 and residues in the active site of Cdk2, we found other interactions that appear play an important role in anchoring and stabilizing the 3_10_-helix in p27 that contain the bulky residues. For example, K33 in Cdk2 — a highly conserved residue (Fig. S4) that extends out from the periphery of Cdk2 active site — is able to hydrogen bond with the backbone carbonyl groups of p27 F87 and R90 (Fig. 3e), and indeed burial of Y88 appears to occur concomitantly with the formation of one or two of these hydrogen bonds (Fig. 3b). Finally, we note that Y74 (which may be phosphorylated after Y88) remains bound even when Y88 is exposed, though we do observe some minor increased dynamics of Y74 once Y88 is exposed (Fig. 3f).

**Figure 3.**
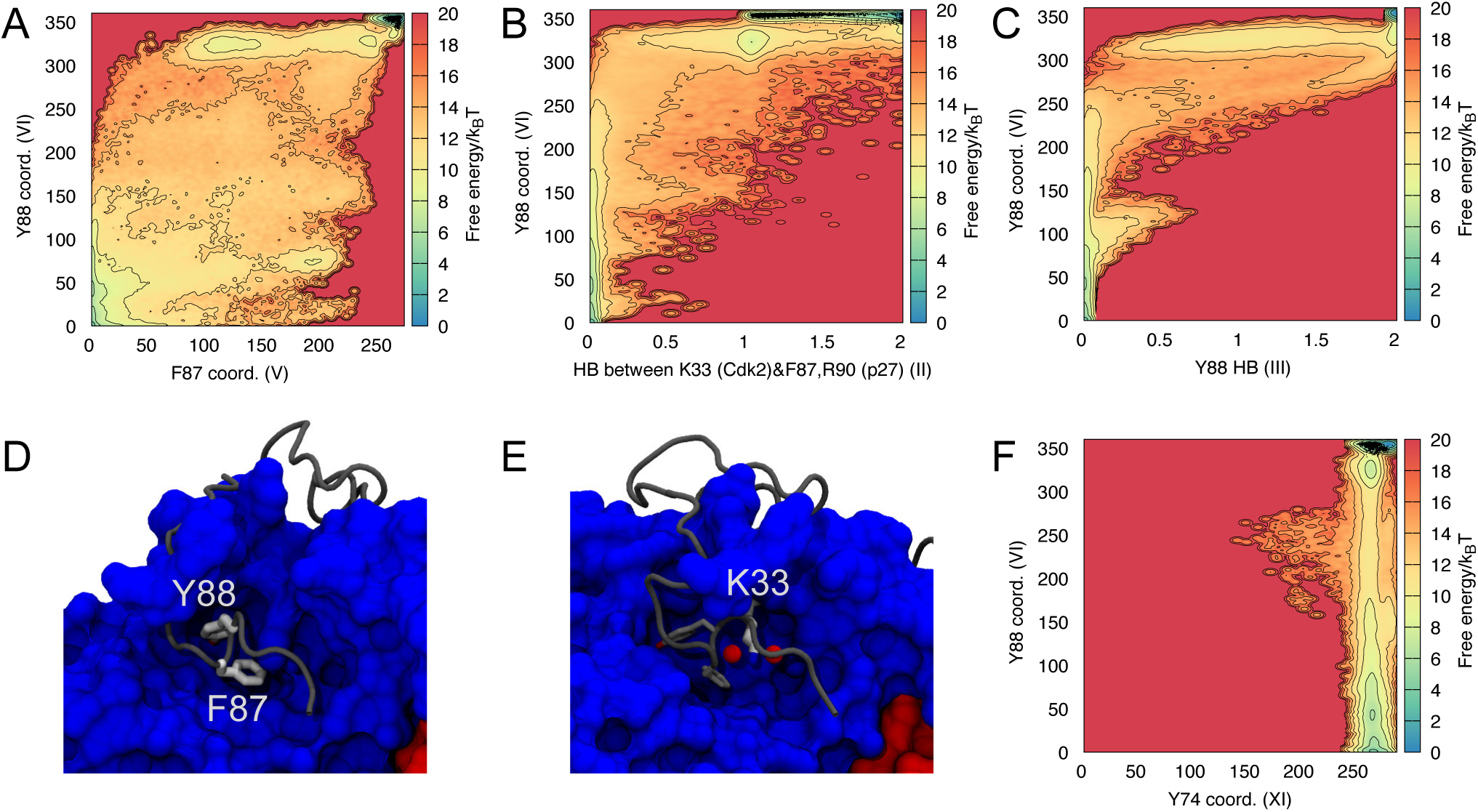
Key interactions governing binding and release of Y88. Selected free-energy profile reconstructions from multiple walkers multiple-replica parallel-bias simulation (Table S1, ref. D) showing: (a) the cooperative binding of F87 and Y88 to the Cdk2 binding site, the stabilization of the bound state through key polar interactions between (b) the side chain of Cdk2 K33 and the backbone of p27, and (c) a hydrogen bond from Y88 itself. Representations of (d) F87 and Y88 in the active site of Cdk2 and (e) Cdk2 K33 and the backbone carbonyl oxygen atoms of p27 F87 and R90. (f) The free-energy profile of CVs capturing contacts between Cdk2 and Y88/Y74, respectively, show that Y74 stays bound in these simulations. Black dots in panels (a), (b), (c) and (f) represent conformations from the unbiased MD simulation starting from the X-ray structure (Table S1, ref. B).

As proposed earlier (***Russo et al., 1996***), several of these interactions may also be important for the binding and stabilization of ATP in the active binary complex (***Brown et al., 1999***) (PDB 1QMZ). Through a direct comparison between the crystal structures of inhibited and active Cdk2/cyclin A, the authors note that p27 mimics ATP in the way it interacts with the active site of Cdk2. F87 and Y88 bind deep in the catalytic cleft, occupying most of the available space in an orientation that traces the position of ATP. In fact, the position and contacts made by Y88 remarkably mimic those of the purine group of ATP. Furthermore, the backbone carbonyl groups of p27 F87 and R90 hydrogen bond with the conserved Cdk2 K33 in similar fashion to the ATP phosphates in the active binary complex (***Russo et al., 1996***).

### Phosphorylation of Y88 leads to a destabilization of the bound state

Having investigated the intrinsic dynamics of the unphosphorylated Cdk2/Cyclin A-bound p27 system, and in particular of Y88, we now focus on the effect of phosphorylation on this assembly. Y88 in p27 is the target of non-receptor tyrosine kinases BCR-ABL (***Grimmler et al., 2007***) and Src (***Chu et al., 2007***), and its phosphorylation represents the first step in a cascade of events that culminate in the full reactivation of Cdk2/Cyclin A and cell cycle progression.

Inspection of the structure of the unphosphorylated ternary complex suggests that the buried conformation of Y88 would be incompatible with Y88-phosphorylated (pY88) p27. Indeed, when we initiated simulations where we had modelled pY88 in the conformation of the stable ternary complex of Y88 (Table S1, ref. F) we find that pY88 rapidly (*<* 100 ns) and spontaneously leaves the binding pocket in Cdk2 (Fig. S5). The observation that pY88 is unstable in the bound configuration is in agreement with the experimentally observed change in the relative populations of the so-called major and minor states of Cdk2/Cyclin A-bound p27 (i.e., buried *vs*. NRTK accessible Y88, respectively) (***Tsytlonok et al., 2019***), and with an earlier computational study (***Rath and Senapati, 2016***).

We observe that when pY88 leaves the ATP-pocket in Cdk2, the conformational changes in p27 are mostly observed around the D2.3 domain, i.e. around residues 83 − 93 (Fig. 2c), similar to the conformational dynamics we observed in the enhanced sampling simulations of p27 with Y88. In other words, the conformational ensemble of pY88 p27 resembles the transiently exposed Y88 state. Of note, the intermolecular *β* strand formed by residues 75 − 79 in p27 and *β*-strand 2 (*β*2) in Cdk2 is for the most part structurally preserved, as are the interactions between the rest of the D2, LH and D1 domains (***Sivakolundu et al., 2005***) of p27 and Cdk2 and cyclin A.

### Phosphorylation of Y88 promotes the exposure of the Y74 phosphorylation site

We proceeded to study the transient exposure of Y74 and how it might be coupled to the phosphorylation status of Y88. We thus performed simulations of (i) unphosphorylated p27 bound to Cdk2/cyclin A and (ii) the same system but with Y88 phosphorylated and thus unbound. In both cases we enhanced sampling using metadynamics by biasing the number of contacts formed by Y74 in p27 (Fig. 4b; Table S3, CV XI) as well as other CVs (Table S3, CVs IX and X). In this way, we aimed to obtain insight into whether the phosphorylation status of Y88 had any effect on the exposure of Y74, and if so by which mechanism. In both the simulation of the unphosphorylated p27 as well as that with pY88, we are able to observe reversible dissociation and rebinding of Y74, both with Y88 in its unphosphorylated state (SI Movies 3 and 4) and with pY88 (SI Movie 5), suggesting that our selection of collective variables captures the main interactions responsible for this process (Fig. 4a).

**Figure 4.**
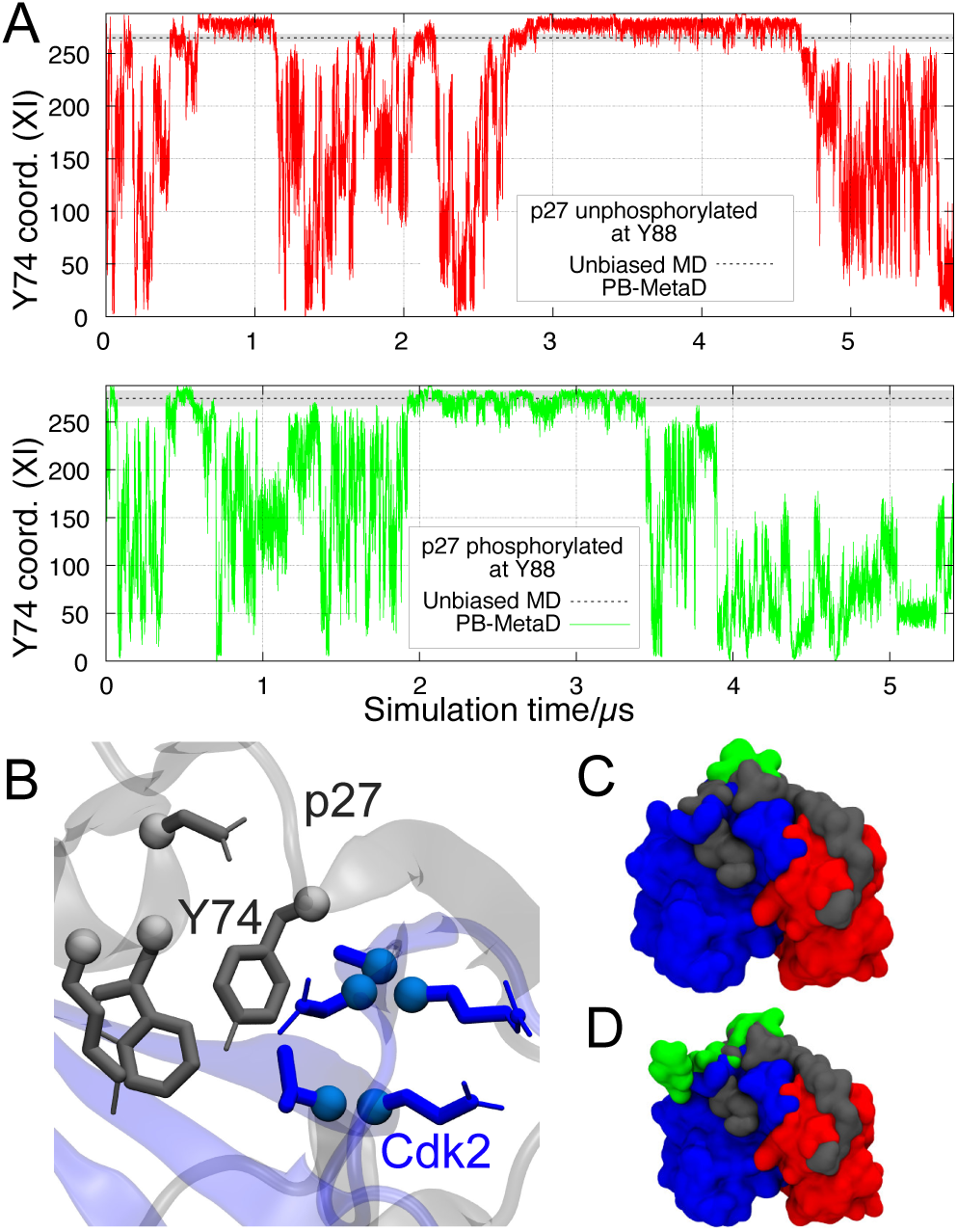
Transient exposure of Y74 in metadynamics simulations. (a) Time series of a CV that quantifies the number of contacts between Y74 and Cdk2 (CV XI, Table S2) in either a parallel bias metadynamics simulation of (upper panel) unphosphorylated p27 Cdk2/cyclin A (Table S1, ref. G) or (lower panel) singly phosphorylated (pY88) p27 (Table S1, ref. H). The average number of contacts calculated from unbiased MD simulations of these systems (Table S1, refs. B and F) are shown with a dashed line. (b) The contacts formed between Y74 (in gray, center) and its neighboring residues in p27 and Cdk2 (gray and blue, respectively) is a good descriptor of the extent to which Y74 is buried in the interface between p27 and Cdk2. (c) An unphosphorylated native-like p27 structure is observed at high coordination numbers (gray), whereas low coordination numbers coincide with the solvent exposure of Y74 (green). (d) Same as in (c) but for the singly phosphorylated system where pY88 is also exposed.

In our simulations, it appears that the phosphorylation state of Y88 affects how easy it is for Y74 to dissociate from its binding pocket (Fig. 5) and also suggest that Y74 is more tightly bound than Y88 (Fig. S6). specifically our simulations suggest that upon phosphorylation of Y88, solvent-exposed Y74 becomes the dominant state (Fig. 5a and 5c), whereas the bound state of Y74 is the preferred state when Y88 is unphosphorylated and bound (Fig. 5a and 5b). A substantial shift in the population of the exposed state Y74 — depending on the phosphorylation status of Y88 — is in agreement with experiments (***Tsytlonok et al., 2019***); however, because of the difficulty in converging the computational free-energy landscapes we defer from making quantitative comparisons. Instead, we proceeded to analyse the structures observed in the simulations to provide insight into how such a population shift might be achieved. We observe that the motions of the intermolecular *β*-sheet formed between the D2.2 domain of p27 and the *β*2 strand in Cdk2 (Fig. 5d) are affected by the phosphorylation status of Y88. Upon phosphorylation, pY88 and the D2.3 domain of p27 are no longer constrained to Cdk2, putting significant strain on this intermolecular *β*-sheet, which then becomes the last point of stable interaction between p27 and Cdk2. As a results this *β*-sheet becomes distorted towards its C-terminal region, enabling an easier tear of its N-terminal contacts.

**Figure 5.**
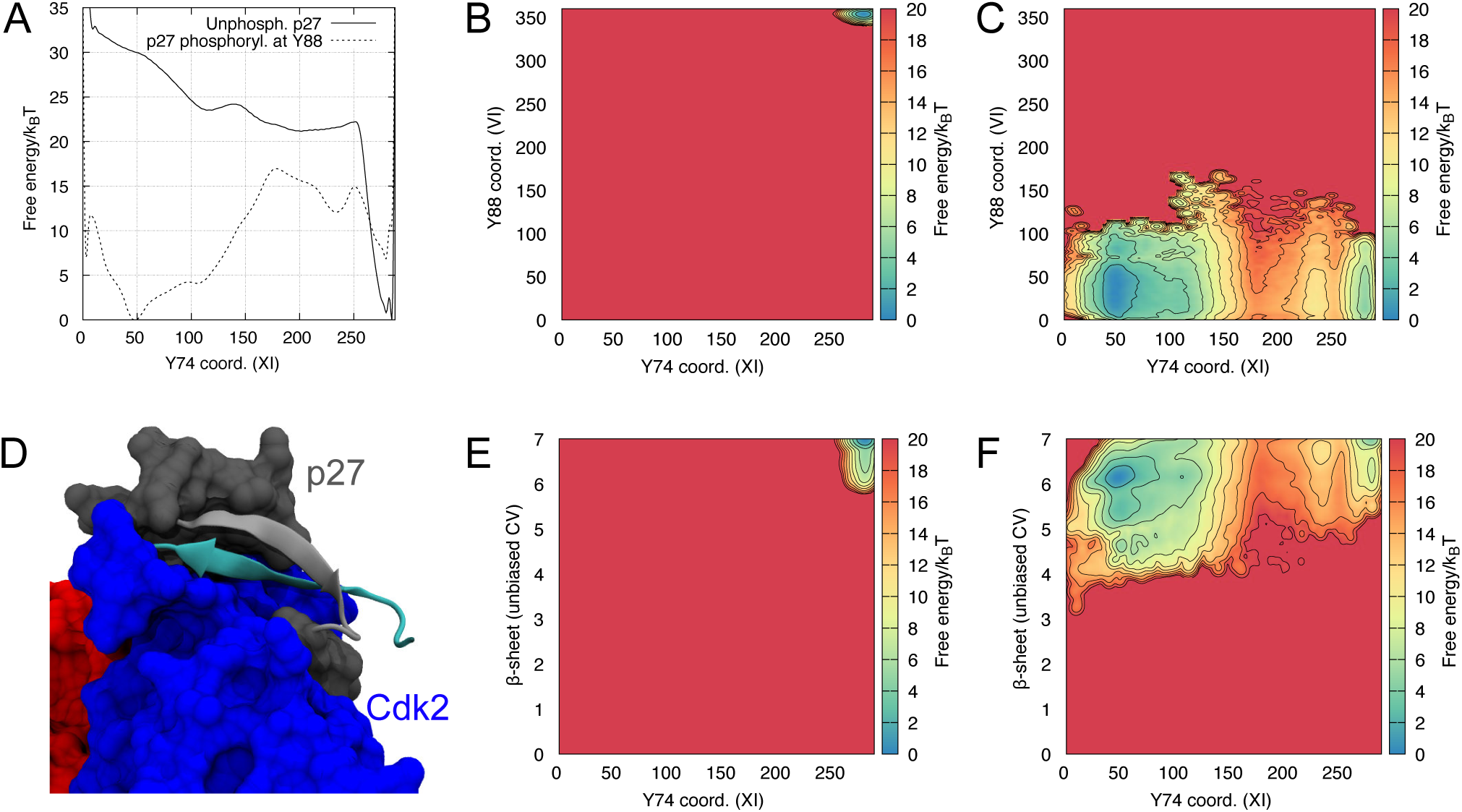
Potential coupling between exposure and phosphorylation of Y88 and Y74. Selected free-energy profiles showing: (a) the equilibrium between buried and solvent exposed Y74 states when Y88 is either phosphorylated or not; the relationship between the degree of exposure of Y74 and either (b) Y88 or (c) pY88; the relationship between the number of contacts in the *β*-sheet formed between p27 and Cdk2 and the degree of exposure of Y74 either with (e) Y88 or (f) pY88. The free-energy profiles are based on the parallel-bias metadynamics simulations (Table S1, refs. G (Y88) and H (pY88)). Panel (d) Illustrates the location of the *β*-sheet formed between p27 and Cdk2 whose number of contacts are used panels (e) and (f).

Because Y74 is located immediately before the start of this *β*-sheet, rupture of contacts on its end will inevitably facilitate its solvent exposure (Fig. 5e-f and SI Movie 5). These observations are also in line with previous simulation results (***Rath and Senapati, 2016***) that noted a relationship between the phosphorylation status of Y88 and the disruption of this intermolecular *β*-sheet. Indeed, in a relatively short (0.5 µs) unbiased MD simulation of doubly phosphorylated p27 bound to Cdk2/Cyclin A (Table S1, ref. I) we observe substantial dynamics of the shared *β*-sheet (SI Movie 6) which would eventually break, leading to the possibility of Cdk2 phosphorylating T187 in p27 in *cis*.

## Conclusion

Protein phosphorylation is a key regulatory mechanism across many biological processes, and phosphophorylation can affect a range of molecular properties. While many phosphorylation events occur in disordered regions of proteins, it can also occur within folded proteins or IDPs that have folded upon binding to their target. The finding that several phosphorylated residues are buried inside a protein or complex in the unphosphorylated state begs the question of how such buried residues become accessible to kinases. Experimental studies of Vav1 (***Li et al., 2008***) and the complex between p27 and Cdk2/Cyclin A (***Tsytlonok et al., 2019***) suggest that protein dynamics may lead to transient exposure of the phosphosite, thus making the residue available for interacting with the kinase. In the case of both Vav1 and the p27 complex, the unphosphorylated, bound state is (auto)inhibited with the relevant tyrosine residues playing a key role in inhibition. This in turn suggests that the equilibrium between a bound and a transiently exposed state would be 1nely tuned because too high a population of the bound state would make phosphorylation exceedingly slow, and a too high population of the exposed state would make inhibition inefficient. In the case of Vav1, mutations that shift the equilibrium between buried and exposed states cause a corresponding change in the rate of phosphorylation (***Li et al., 2008***), suggesting a conformational selection mechanism for the kinase rather than an active role in helping expose the phosphosite.

We have here studied the intrinsic dynamic motions of p27 when bound to Cdk2/Cyclin A that are relevant for phosphorylation of Y88 and Y74. In line with recent experiments, which were published after this work was initiated, we find that both phosphorylation sites can form transiently exposed states, and that these are high in free energy. Thus, transient exposure of Y88 is a rare event so that Cdk2 is almost fully inhibited. Once Y88 is phosphorylated, our simulations suggest that Y74 becomes more easily exposed, and that the shared *β*-sheet between Cdk2 and Cyclin A is important for mediating this coupling. Our work was made possible by using enhanced sampling simulations and CVs selected based on the crystal structure of the complex, and the recent development of force fields that are accurate for both folded and disordered proteins (***Robustelli et al., 2018***). Our difficulties in obtaining converged free energy landscape complicated direct comparisons to NMR and single molecule fluorescence experiments (***Tsytlonok et al., 2019***). Future studies could benefit from a more direct integration of experiments and simulations, though are in part hampered by problems with forward models for NMR chemical shifts and difficulties in incorporating kinetic data into such models (***Orioli et al., 2020***). We envisage that simulations of this kind may be used more generally to study the dynamic motions that we suggest might underlie phosphorylation of buried residues in many other proteins.

## Methods

### Protein structure

We used the crystal structure of the ternary complex Cdk2/Cyclin A/p27 (PDB (***Berman et al., 2000***) entry 1JSU (***Russo et al., 1996***)) as starting point for our simulations. In this structure, portions of the N- and C-termini of Cdk2 and p27, respectively, were not resolved, and we therefore capped these with acetyl and N-methyl groups. We mutated Y89 in p27 to a phenylalanine to match the construct used in previous NMR and fluorescence experiments (***Tsytlonok et al., 2019***). We set the residue protonation to the most probable state at pH 7.5 and used the GROMACS pdb2gmx tool (***Abraham et al., 2015***) to determine histidine tautomerism based on each residue’s environment. In some of our simulations (see Table S1 for an overview of the simulations that we performed), p27 was phosphorylated at either Y88 or Y88 and Y74 with individual net charge −2 *e*.

### Molecular dynamics

We used GROMACS (version 2016.4) (***Abraham et al., 2015***) for our MD simulations, in most cases using the AMBER ff99SB-*disp* force field (***Robustelli et al., 2018***), though we also used CHARMM36m (***Huang et al., 2017***) with the CHARMM-modified TIP3P water model (***MacKerell Jr et al., 1998***) for certain control simulations. AMBER force field parameters for the phosphotyrosine were adapted from previous parameterizations (***Homeyer et al., 2006***; ***Steinbrecher et al., 2012***). The simulated systems consist of the ternary complex in a rhombic dodecahedral simulation box (*d* = 10.5 nm), filled with pre-equilibrated water molecules (at 300 K and 1 bar) and the necessary number of Na^+^ and/or Cl^−^ atoms required to neutralize the system. Periodic boundary conditions were employed in all directions.

We used the leapfrog algorithm (***Hockney and Eastwood, 1988***) with a time step of 2 fs. Lennard-Jones and short-range electrostatic interactions were calculated using Verlet lists with cutoffs of 1.0 and 1.2 nm, for AMBER and CHARMM force fields, respectively. Lennard-Jones interactions were switched off smoothly (force switch) between 1.0 nm and the aforementioned 1.2 nm cutoff for simulations using CHARMM36m. Long-range dispersion corrections were applied to the energy and pressure when using AMBER ff99SB-*disp*. The particle-mesh Ewald method (***Darden et al., 1993***) was used to treat long-range electrostatic interactions, using a grid spacing of 0.12 nm and cubic interpolation. Solvent and solute were coupled to temperature baths at 300 K, using the velocity rescaling thermostat (***Bussi et al., 2007***) with a relaxation time of 0.1 ps. The Parrinello-Rahman barostat (***Parrinello and Rahman, 1981; Nosé and Klein, 1983***) was set to 1 bar, with a relaxation time of 2 ps and isothermal compressibility of 4.5 × 10^−5^ bar^−1^. Bond lengths were constrained using the LINCS algorithm (all bonds for AMBER ff99SB-*disp* and bonds with hydrogen atoms for CHARMM36m) (***Hess et al., 1997***). We used a steepest descent algorithm for minimization. Initiations were performed in a two-step scheme for a total of 2 ns, using position restraints of 1000 kJ mol^−1^ nm^−2^ on all protein heavy atoms. The first step was performed under the canonical ensemble to stabilize the temperature, and the second step was performed under the same ensemble as the production simulations, i.e., the isobaric-isothermal ensemble. Protein coordinates were stored every 10 ps. Simulation lengths are reported in Table S1 and range from 0.1 µs–5.7 µs.

### Metadynamics simulations

We used the PLUMED plug-in (version 2.4.1) (***Sutto et al., 2012***) to perform (well-tempered) meta-dynamics simulations (***Laio and Parrinello, 2002; Pfaendtner and Bonomi, 2015; Barducci et al., 2008***). specific parameters and collective variables (CVs) that we used are presented in the Supporting Information (Tables S2 and S3), and here we just describe the most central ones. The placement of Y88 in its binding pocket is quantified via the calculation of the coordination number (number of contacts) between the atoms in Y88 and a number of heavy atoms in Cdk2 that are within 6Å in the crystal structure (CV: VI). We calculated the number of contacts for F87 and Y74 in a similar way (CVs V and XI, respectively). In addition to these CVs that quantify how well these three aromatic residues dock into their pockets, we used additional CVs to capture key hydrogen bonding interactions between p27 and Cdk2 (CV II) and a hydrogen bond between the Y88 side chain and Cdk2 (CV III).

When explicitly stated, multiple replicas of the same system — using the same set of collective variables and parameters, sharing and contributing to the same overall history dependent bias potential — were run in parallel, in what is often referred to as multiple walkers metadynamics (***Raiteri et al., 2006***), and was combined with parallel bias metadynamics simulations (***Pfaendtner and Bonomi, 2015***). We updated the bias potential every 2 ps, with an initial bias deposition of 1 kcal/mol and a ‘bias factor’ of 12.5 (later on increased to 15 in some simulations). Protein and full system coordinates were stored every 2 ps and 1 ns, respectively. All other parameters are similar to those described above for the unbiased MD simulations. Further details can be found in Table S1.

### Analysis

We used PLUMED (version 2.4.1) (***Sutto et al., 2012***) and *in-house* tools to analyse the simulations. One-dimensional free-energy profiles were constructed by summing all deposited Gaussians during the metadynamics simulation (PLUMED sum_hills tool). The weights needed to construct the two dimensional free-energy surfaces were estimated (***Pfaendtner and Bonomi, 2015***) after discarding the initial 200 ns from the trajectories of each metadynamics simulation for equilibration (Fig. S7). All plots were generated using GNUPLOT (***Williams et al., 2019***), and protein images were rendered using VMD (***Humphrey et al., 1996***) and the Tachyon ray tracing software (***Stone, 2014***).

## Author Contributions

K.L.-L. conceived the idea; J.H. performed and analysed simulations; J.H. and K.L-L. designed the research, analysed the data and wrote the manuscript

## Acknowledgments

This work was supported by Independent Research Fund Denmark (grant number 6108-00471) and the Lundbeck Foundation BRAINSTRUC initiative in structural biology. We thank Richard Kriwacki and Claus Seidel for fruitful discussions, and Sandro Bottaro, Daniele Granata, Simone Orioli, Nour Saleh and Yong Wang and other members of the Lindorff-Larsen group for help and discussions.

