## Supporting information for "Protein dynamics enables phosphorylation of buried residues in Cdk2/Cyclin A-bound p27"

### List of Simulations

Table 1: List of simulations performed and analyzed in this work.

| Ref. | Start configuration | Force field | Water model[4] | Type[5] | No. walkers[6] | Sim. length[6](μs) |
|---|---|---|---|---|---|---|
| A | Native[1] | CHARMM36m | TIP3P | MD | – | 2 |
| B | Native[1] | AMBER ff99SB-*disp* | TIP4P-D | MD | – | 0.5 |
| C | Native[1] | AMBER ff99SB-*disp* | TIP4P-D | PB-MetaD | 1 | 5 |
| D | Native[1] | AMBER ff99SB-*disp* | TIP4P-D | PB-MetaD | 8 | 2 |
| E | Y88 intermediate state[1] | AMBER ff99SB-*disp* | TIP4P-D | MD | – | 0.1 |
| F | Y88 bound & phosp.[2] | AMBER ff99SB-*disp* | TIP4P-D | MD | – | 0.5 |
| G | Native[1] | AMBER ff99SB-*disp* | TIP4P-D | PB-MetaD | 1 | 5.7 |
| H | Y88 unbound & phosp.[2] | AMBER ff99SB-*disp* | TIP4P-D | PB-MetaD | 1 | 5.4 |
| I | Y88/Y74 unbound & phosp.[3] | AMBER ff99SB-*disp* | TIP4P-D | MD | – | 0.5 |

[1] PDB 1JSU, TPO160 → T160, ACE-cap on Cdk2, NME-cap on p27.
[2] Same as in [1], but with p27 phosphorylated at Y88.
[3] Same as in [1], but with p27 doubly phosphorylated at Y88 and Y74.
[4] For the CHARMM simulation we used CHARMM-modified TIP3P, and for the AMBER simulations we used the water model native to ff99SB-*disp* which is similar to, but not identical with, the TIP4P-D model.
[5] MD here refers to unbiased MD and PB-MetaD to parallel-bias metadynamics simulations
[6] For parallel-bias metadynamics simulations we report the number of walkers used and the length of the simulation for each walker.

### Metadynamics

#### Collective variables (CVs)

All CVs that we used involve the number of contacts formed between two groups of atoms $\mathcal{A}$ and $\mathcal{B}$, commonly referred to as the coordination number, here defined as:

$$coord_{\mathcal{AB}} = \sum_{i\in\mathcal{A}}\sum_{j\in\mathcal{B}} \frac{1-\left(\frac{r_{ij}-d_0}{r_0}\right)^6}{1-\left(\frac{r_{ij}-d_0}{r_0}\right)^{12}} \ , \tag{1}$$

where $r_{ij}$ is the instantaneous distance between atoms $i$ and $j$, while $r_0$ and $d_0$ are intrinsic switching function parameters. For each contact, when $r$ becomes much smaller than $d_0$ then $coord_{ij}$ goes towards unity, and when $r$ becomes greater than $d_0$ it decays smoothly to 0. At $r = d_0$, $coord_{ij} = 0.5$.

Atom selections, switching parameters, Gaussian width ($\sigma$ ) and CV space for simulations C & D and G & H (Table 1) are presented in Tables 2 and 3, respectively.

Table 2: Collective variables used to bias simulations C & D (Table 1).

| CV | Group $\mathcal{A}$ | Group $\mathcal{B}$ | $r_0$ (Å) | $d_0$ (Å) | $\sigma$ | Grid space |
|---|---|---|---|---|---|---|
| I | p27 K81 main-chain N atom | Cdk2 G16 main-chain O atom | 3.5 | 3.5 | 0.05 | $0-1$ |
| II | p27 F87 & R90 main-chain O atoms | Cdk2 K33 side-chain N atom | 3.5 | 3.5 | 0.1 | $0-2$ |
| III | p27 Y88 side-chain O atom | Cdk2 E81 main-chain O & L83 main-chain N atoms | 3.5 | 3.5 | 0.1 | $0-2$ |
| IV | p27 R90 side-chain N atoms | Cdk2 Q131 main-chain O, N132 & D145 side-chain O atoms | 3.5 | 3.5 | 0.4 | $0-8$ |
| V | p27 F87 $C_\alpha$ & side-chain C atoms | $C_\alpha$ & side-chain C atoms of Cdk2 residues within 6 Å of group $\mathcal{A}$ | 6 | 6 | 1.5 | $0-272$ |
| VI | p27 Y88 $C_\alpha$ & side-chain C atoms | $C_\alpha$ & side-chain C atoms of Cdk2 residues within 6 Å of group $\mathcal{A}$ | 6 | 6 | 1.5 | $0-360$ |
| VII | p27 Y89F $C_\alpha$ & side-chain C atoms | $C_\alpha$ & side-chain C atoms of p27/Cdk2 residues within 6 Å of group $\mathcal{A}$ | 6 | 6 | 1.5 | $0-104$ |
| VIII | Cdk2 G13−V18 $C_\alpha$ & $C_\beta$ atoms | Cdk2 $C_\alpha$ & $C_\beta$ atoms within 7 Å of p27 F87 & Y88 | 18 | 0 | 5 | $0-260$ |

Table 3: Collective variables used to bias simulations G & H (Table 1).

| CV | Group $\mathcal{A}$ | Group $\mathcal{B}$ | $r_0$ (Å) | $d_0$ (Å) | $\sigma$ | Grid space |
|---|---|---|---|---|---|---|
| IX | p27 E75 main-chain N atom | Cdk2 R22 main-chain O atom | 3.5 | 3.5 | 0.05 | $0-1$ |
| X | p27 Y74 main-chain N & O atoms | Cdk2 N61 side-chain O & N atoms | 3.5 | 3.5 | 0.1 | $0-2$[1] |
| XI | p27 Y74 $C_\alpha$ & side-chain C atoms | $C_\alpha$ & side-chain C atoms of p27/Cdk2 residue within 6 Å of group $\mathcal{A}$[2] | 6 | 6 | 2.5 | $0-288$ |

[1] PLUMED keyword PAIR is used here to enforce the pairing of the 1st element of group $\mathcal{A}$ with 1st element of group $\mathcal{B}$, and so forth, avoiding the inclusion of electrostatically unfavourable contacts (e.g. carbonyl-carbonyl oxygen interactions).

[2] p27 Y74 intrachain contacts are only allowed when they are 5 or more residues apart.

### Supporting figures

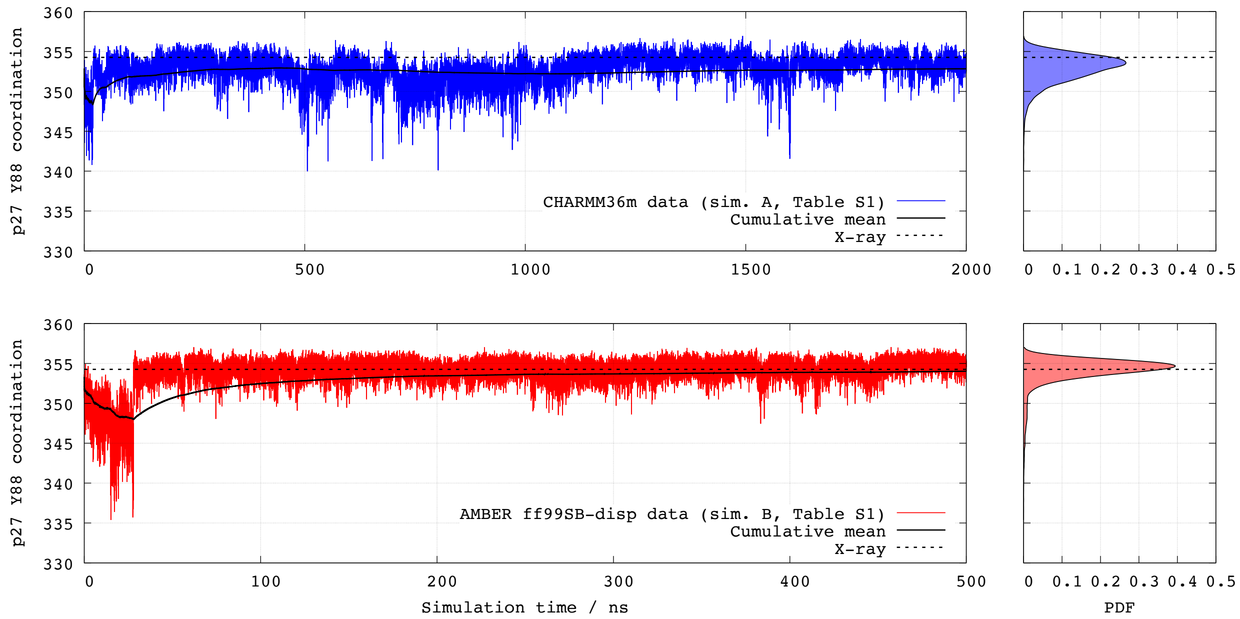


Figure 1: Analysis of the coordination number collective variable for Y88 (CV VI, Table S2) in an unbiased MD simulation. The plots show the time series, cumulative mean and probability density functions (PDF) computed from unbiased MD simulations of PDB 1JSU using the CHARMM36m and AMBER ff99SB-*disp* force fields (upper and lower panel, respectively). Similar results were obtained for Y74.

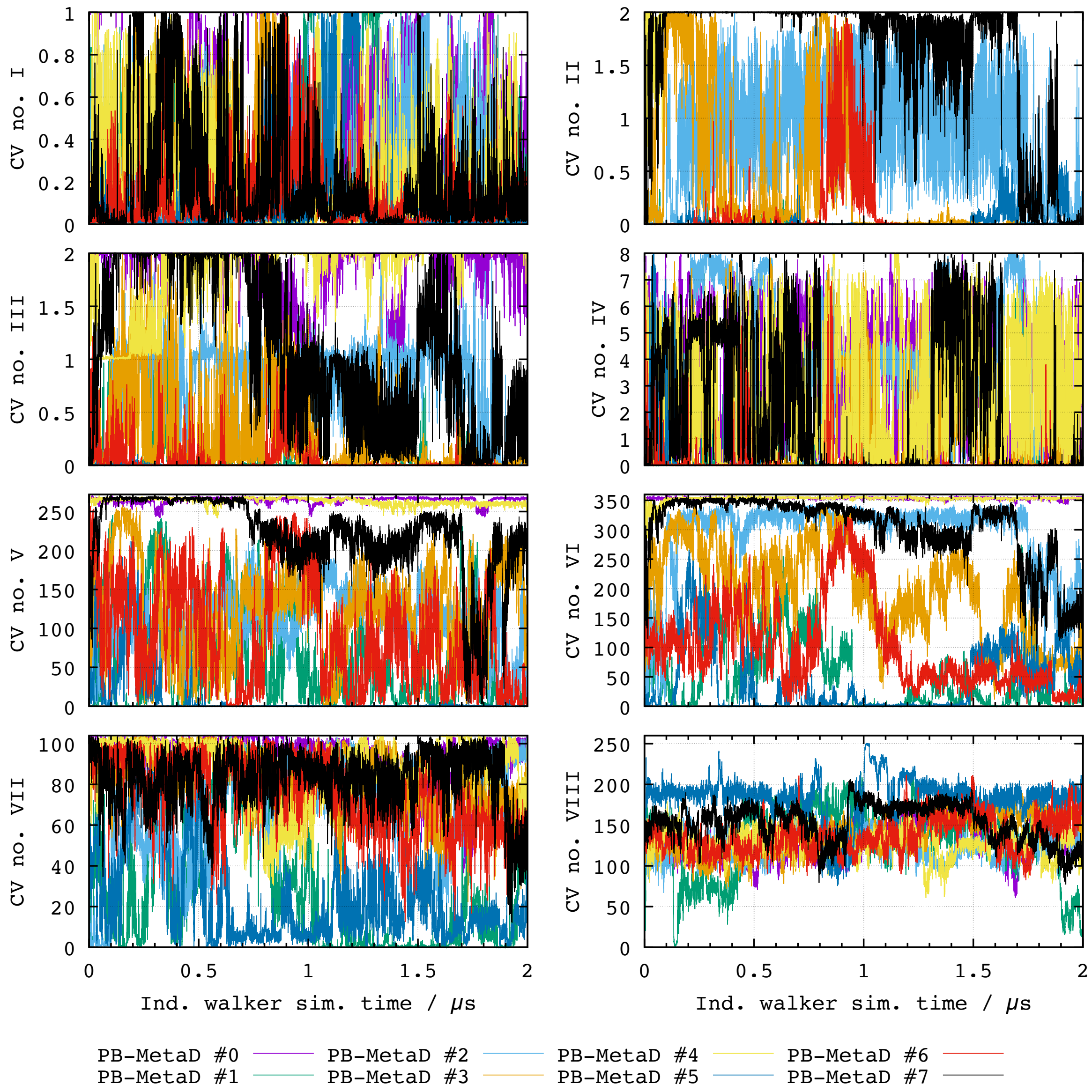


Figure 2: Time series for each biased CV (see Table 2) in the multiple walkers parallel-bias metadynamics simulation of the unphosphorylated Cdk2/Cyclin A-bound p27 system (Table 1, Ref. D). The results are shown for all eight individual walkers.

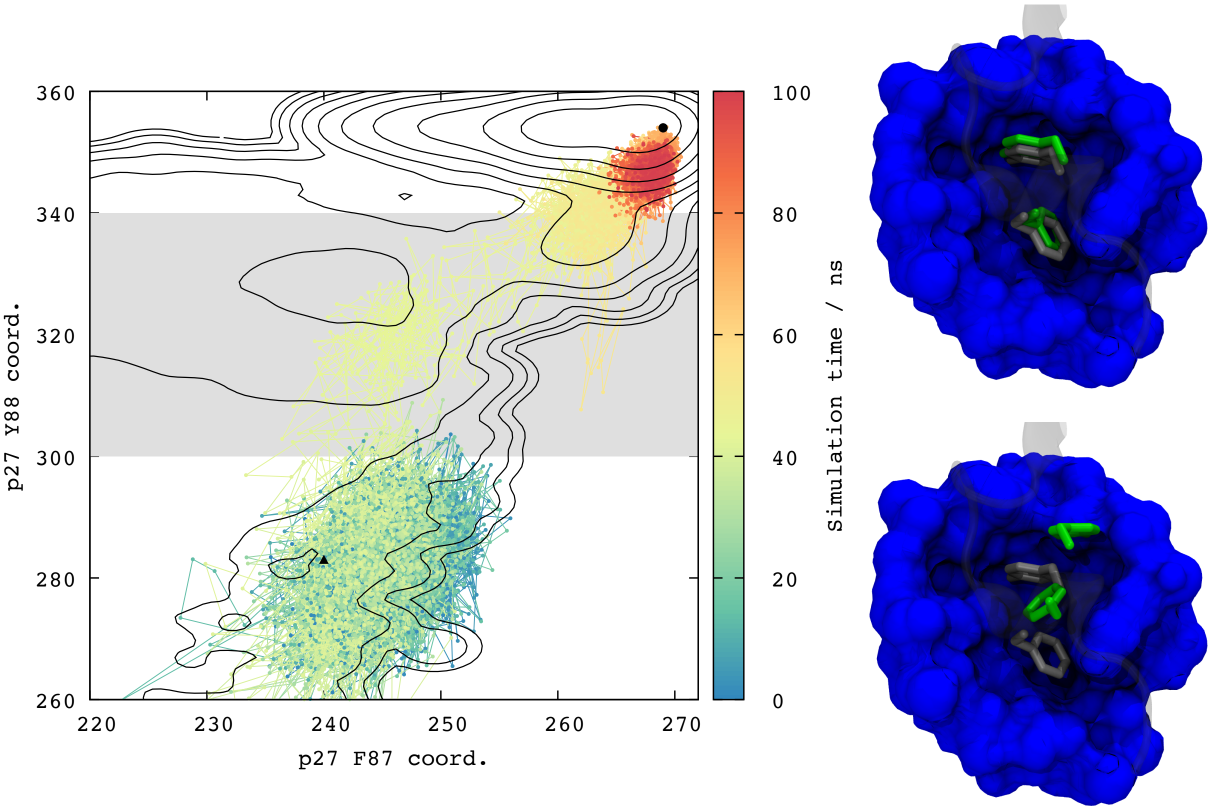


Figure 3: Spontaneous binding of Y88 to the ATP binding state in unbiased simulations initiated from an intermediate state. (Left) Time progression of a MD simulation (Table S1, Ref. E) represented along a free energy profile spanned by the coordination numbers of F87 and Y88 (CVs VI and V, respectively; Table S2). The simulation was initiated from an intermediate state with Y88 partially buried and solvent exposed (black triangle) and spontaneously progressed towards the global minimum region, corresponding to a native-like placement of F87 and Y88 deeply within Cdk2 active site (black circle). The gray shaded area represents the rarely traversed region in the multiple walkers PB-MetaD simulation (Table S1, Ref. D). This simulation crosses this region spontaneously, following the lowest energy path delimitated by the black contour lines (taken from Fig. 3A) with the colour indicating the simulation time. (Right) Final (top) and starting (bottom) states, corresponding to the black triangle and circle in (a), respectively. The location of F87 and Y88 in the crystal structure are represented in gray and the conformation from MD are represented in green. Part of the active site of Cdk2 is represented in blue and p27 was made transparent. Structure alignments were performed using the backbone of Cdk2 and cyclin A only.

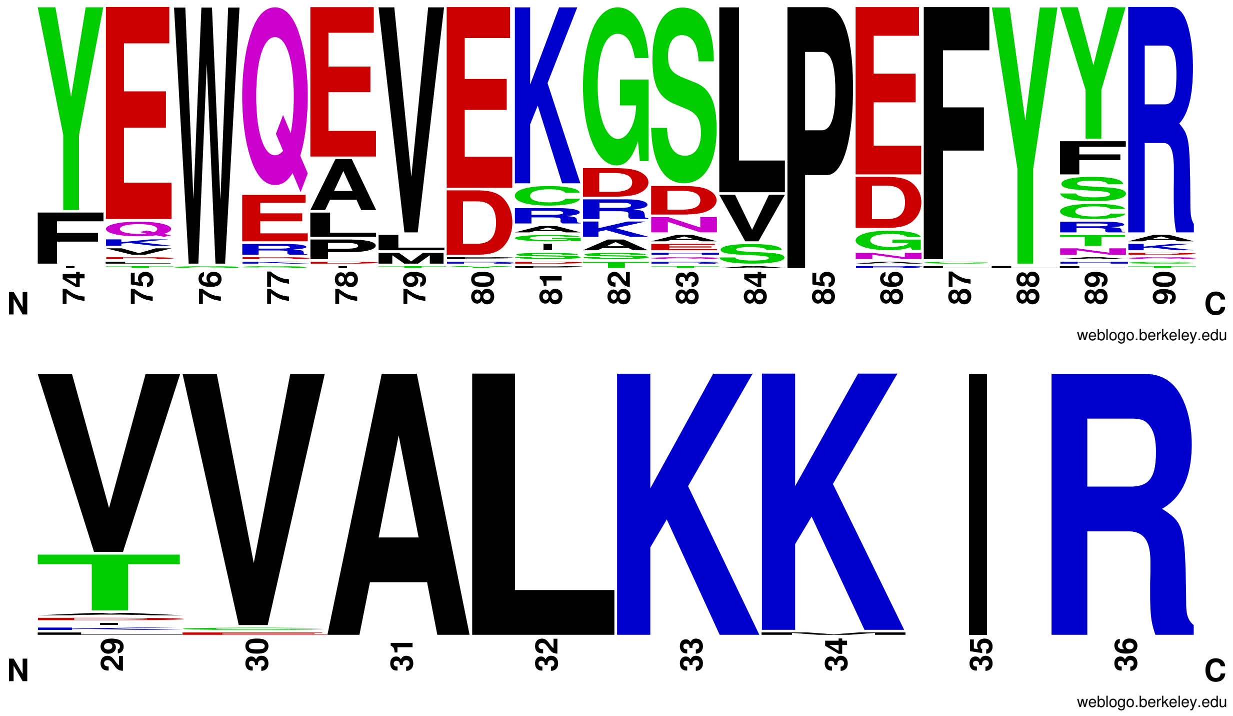


Figure 4: Logo plots highlighting the sequence conservation of (a) part of the p27 D2 domain and (b) the third $\beta$ strand in Cdk2, using 65 orthologues of the human CDKN1B and CDK2 genes, respectively.[1] p27 F87 and Y88 are highly conserved throughout all available species, as is K33 in Cdk2. p27 Y74 is less well conserved, often being substituted with a phenylalanine residue. Images were generated using the weblogo server.[2]

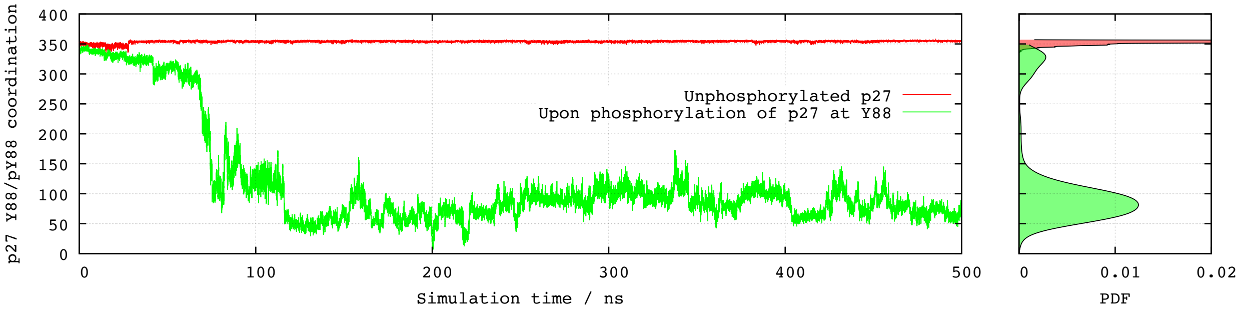


Figure 5: Unbiased MD simulation of Y88-phosphorylated p27 initiated from the bound state. The figure shows the Y88 coordination number collective variable (CV VI, Table S2); (left) time series and (right) probability density functions (PDF) computed from an unbiased MD simulation initiated from the bound state with either Y88 (red; Table S1, ref. B) or pY88 (green, Table S1, ref. F).

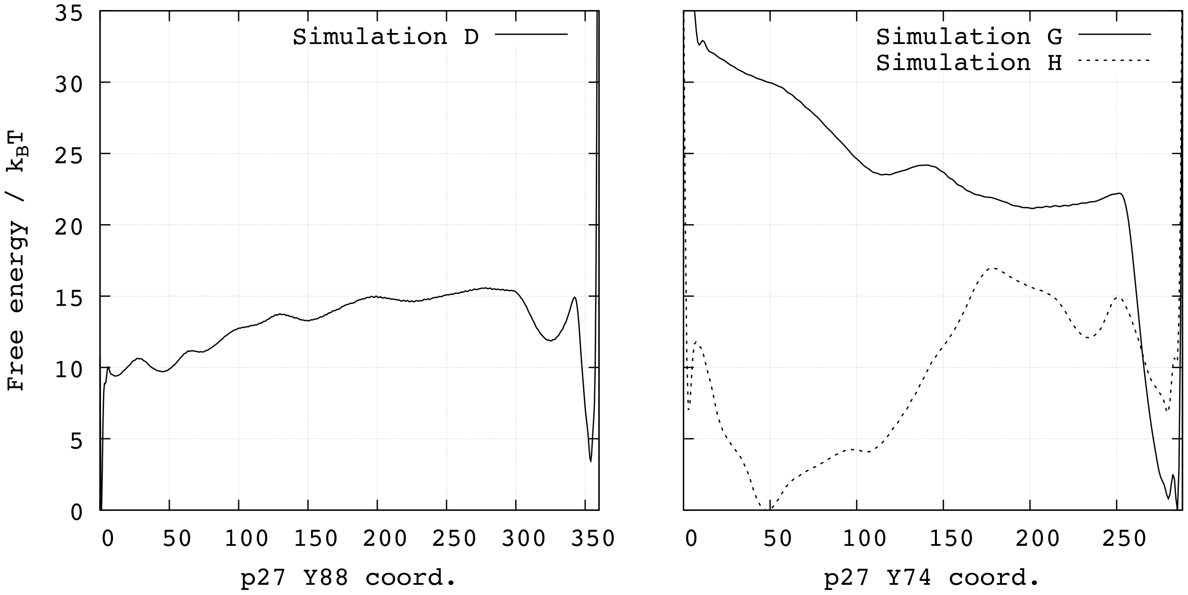


Figure 6: Comparing free energy profiles of exposing Y88 (left) and Y74 (right). The left panel corresponds to the 1-dimensional free energy profile of from the PB-MetaD simulation (Table S1, ref. D). The right panel (identical to Fig. 5A) compares the free energies of exposing Y74 both when Y88 is unphosphorylated (Table S1, ref. G) and phosphorylated (Table S1, ref. H).

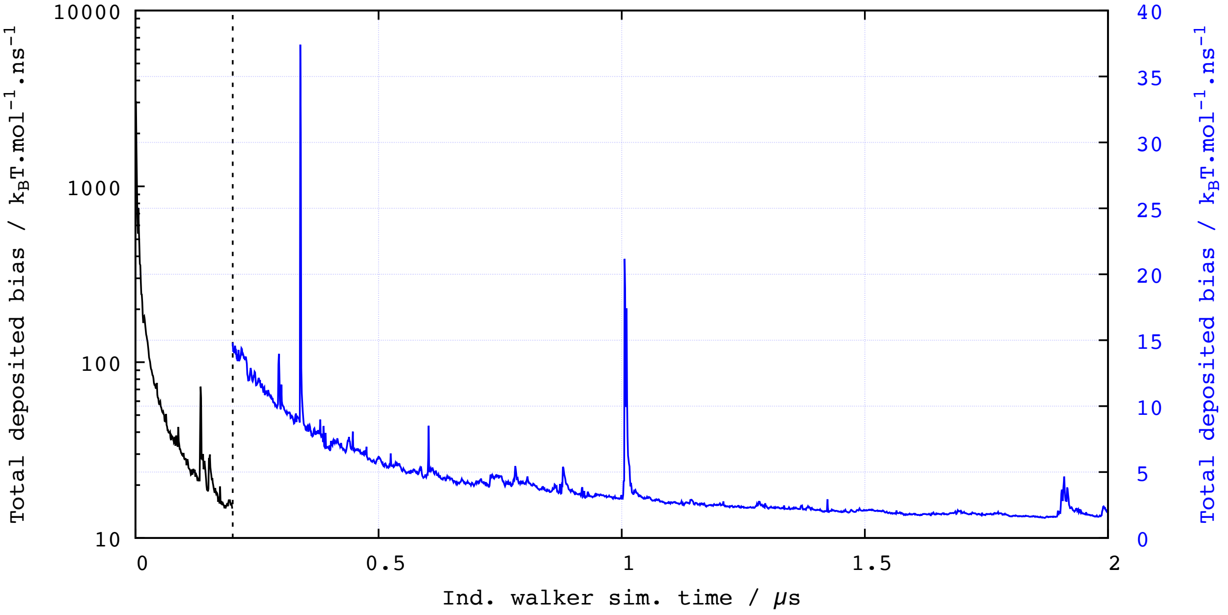


Figure 7: Bias-deposition and equilibration. In parallel bias metadynamics, the construction of 2D or higher dimensional free energy profiles relies on the approximation that the bias potential becomes quasi-static in the long-time limit. In order to minimize the effects of ignoring the time dependence of this potential, we discard the initial 200 ns of the trajectories of all metadynamics simulations, as this is the period where large bias deposition occurs (black line, left y-axis). Throughout the remainder of these simulations, much less bias is added (blue line, right y-axis). This particular plot refers to the multiple walker simulation Ref. D (Table S1). All other metadynamics simulations display a similar profile.
